# Functional specialization of LC3B and LC3C in autophagosome biogenesis and cargo degradation: relevance of the N-terminal region

**DOI:** 10.64898/2026.11.12.766255

**Authors:** Uxue Ballesteros, David Bermejillo, Camila C. Aguirre, Alicia Alonso, L. Ruth Montes, Asier Etxaniz

**Author notes:** Corresponding authors: Asier Etxaniz and L. Ruth Montes. These authors contributed equally to this work (Uxue Ballesteros, David Bermejillo).

## Abstract

ATG8 proteins are key regulators of autophagosome biogenesis and maturation. Despite extensive research on human ATG8 family members, the functional differences between individual proteins remain poorly understood. Among them, LC3C is one of the least characterized members. However, its involvement in a growing number of cellular processes has generated increasing interest in understanding its specific functions. Here, we investigated the functional specialization of LC3C using HeLa ATG8 Hexa-knockout cells individually reconstituted with GFP-LC3C or GFP-LC3B. This experimental system allowed us to examine the specific properties of each protein independently of other ATG8 family members. Under starvation conditions, LC3C formed more numerous and larger autophagic vesicles than LC3B, while both proteins sustained autophagic flux. Furthermore, proteinase K protection assays confirmed that both proteins could promote autophagosome closure. The two proteins also differed in their ability to promote the autophagic degradation of specific cargo receptors, with LC3B being more efficient mediating p62 degradation and LC3C in NDP52 degradation. Confocal microscopy revealed that LC3B and LC3C predominantly localized to distinct autophagic vesicles, suggesting their association with different autophagosome subpopulations. Finally, analysis of chimeras generated by exchanging regions between LC3B and LC3C revealed that the N-terminal region of LC3C determines autophagic vesicle size, whereas its ubiquitin-like core influences intracellular localization. Together, these results suggest that LC3C contributes to determining specific autophagosome properties, such as their size and intracellular localization, expanding our understanding of how individual ATG8 proteins regulate the autophagic process.

## INTRODUCTION

Macroautophagy (hereafter autophagy) is a conserved catabolic process through which intracellular material, including damaged organelles, protein aggregates, and non-specific cytoplasmic components, is delivered to lysosomes for degradation and recycling [1]. Given the essential role of autophagy in cellular homeostasis, autophagy dysfunction has been associated with a wide variety of diseases, including neurodegenerative disorders and aging-related alterations [2]. During autophagy, a membrane structure called the phagophore is formed and expands to engulf intracellular material before closing to form a double-membrane vesicle called the autophagosome (AP), which then fuses with lysosomes for cargo degradation [3,4]. A large number of proteins, known as ATG (AuTophaGy-related) proteins, have been described to participate in AP biogenesis, among which the ATG8 protein family plays a crucial role at several stages of this process [5]. In humans, the ATG8 protein family comprises at least six well-characterized orthologs of the yeast Atg8 protein, divided into two subfamilies: LC3, including LC3A, LC3B and LC3C, and GABARAP, comprising GABARAP, GABARAPL1 and GABARAPL2. While yeast contains a single Atg8 protein, the expansion of this family in higher eukaryotes suggests that individual ATG8 proteins may have evolved to perform specialized functions [5]. To participate in the autophagic process, ATG8 proteins must be covalently conjugated to phosphatidylethanolamine (PE) in the autophagosomal membrane through the coordinated action of two ubiquitin-like conjugation systems [6]. Once anchored to the autophagosomal membrane, ATG8 proteins participate in different stages of the autophagic process, including cargo recognition, phagophore expansion, autophagosome closure and autophagosome-lysosome fusion [7]. Previous studies using HeLa cells lacking all six canonical ATG8 proteins (HeLa HexaKO cells) have shown severe defects in autophagosome biogenesis and maturation, including a reduction in the number and size of autophagosomes, as well as impaired autophagic degradation [8,9]. Despite their established importance in the autophagic process, the specific functions of individual ATG8 family members remain poorly understood. Understanding this functional specialization is therefore essential to elucidate the molecular mechanisms underlying autophagy and its involvement in pathological conditions.

Among members of the LC3 subfamily, LC3B is the most extensively studied and is widely used as an autophagosomal marker, whereas LC3C is the most divergent in terms of amino acid sequence and structure and remains comparatively poorly characterized [10]. The limited characterization of LC3C may be partly explained by its relatively low endogenous expression in commonly used cell models, difficulties in its detection with specific antibodies, and the predominant use of LC3B as a representative of the LC3 subfamily in autophagy studies. LC3C exhibits greater sequence divergence from LC3B (approximately 50% identity) than LC3A does (approximately 80% identity), with differences in its N-terminal region, ubiquitin-like core, and C-terminal extension [11]. In particular, LC3C contains a disordered N-terminal segment corresponding to α-helix 1 in the other LC3 subfamily members [12], as well as specific amino acid substitutions within its LIR-docking site (LDS) that contribute to selective recognition of interacting proteins [13]. In recent years, several studies have reported the involvement of LC3C in different forms of selective autophagy, including xenophagy [14], mitophagy [15], and aggrephagy [16]. In addition, LC3C has been implicated in other cellular processes, such as antiviral defense [17], ER protein export [18], endocytosis [19], exocytosis [20], vesicular trafficking [21], and the regulation of tumor growth [22]. However, the molecular mechanisms underlying the functional specialization of LC3C remain poorly understood.

Among the different functions attributed to ATG8 proteins, their involvement in phagophore expansion remains one of the least understood. Phagophore expansion is primarily driven by the direct transfer of lipids from the endoplasmic reticulum mediated by ATG2 [23], although several studies have suggested that ATG8 proteins may also contribute to this process through a complementary mechanism involving vesicle fusion [24,25]. Our laboratory has investigated the potential contribution of ATG8 proteins to phagophore expansion using an *in vitro* approach with purified proteins and model membranes [26–28], with the hypothesis that these activities may be related to the cellular functions of the proteins [29]. In a recent study, our laboratory showed that LC3C exhibits higher vesicle tethering and membrane fusion activities than LC3B using purified proteins and model membranes. Furthermore, experiments using chimeric proteins identified the N-terminal region of LC3C as responsible for these enhanced activities [30]. In this work, we aimed to determine whether these findings could be extended to a cellular model, to better understand the functional specialization of LC3C.

The study of the specific functions of individual ATG8 family members is complicated due to the functional redundancy among these proteins. ATG8 Hexa-knockout (HexaKO) cells, lacking the six major human ATG8 family members, have been previously used to investigate the functions of ATG8 family proteins [8]. Based on this cellular model, we generated stable cell lines expressing a single GFP-tagged ATG8 family member. This approach allows us to investigate the specific properties of individual ATG8 proteins in the absence of the other family members. In the present study, we used this cellular model to investigate the functional specialization of LC3C in comparison with LC3B. We aimed to characterize the specific properties of autophagosomes formed in the presence of LC3C as the sole ATG8 family member, including their size, intracellular localization, and functionality, and to determine to what extent LC3C alone is sufficient to restore autophagic function. Specifically, we investigated whether the differences in membrane tethering and fusion activities previously observed between these proteins *in vitro* are reflected in their cellular functions. Our findings contribute to a better understanding of the specific functions of LC3C, one of the least characterized members of the ATG8 family.

## MATERIALS AND METHODS

### Materials

Dulbecco’s modified Eagle’s medium (DMEM, D5796), penicillin and streptomycin (P4333-100mL), Luminata Forte Western HRP Substrate (WBLUF0500), Bafilomicin A1 (SML1661.1ML), proteinase K (3115836001) were purchased from Merck (Darmstadt, Germany). Puromycin dihydrochloride from *Streptomyces alboniger* (p388-10MG), RIPA buffer (R0278-50mL), Bovine Serum Albumin (BSA) (A7906) and Mowiol 4-88 (81381-50G) were purchased from Sigma-aldrich (Vienna, Austria). Anti-GFP (mouse, sc-9996), anti-p62 (mouse, sc-28359), anti-vinculin (mouse, sc-73614), anti-mouse IgG-HRP (sc-525409), anti-rabbit IgG-HRP (sc-2352), and Paraformaldehide %4 in PBS (sc-281692) were provided by Santa cruz Biotechnology (Heiderlberg, Germany). Anti-ATG9A (ab108338) and anti-NDP52 (ab68588) were provided by Abcam (Cambridge, UK). Gibco fetal bovine serum (10270106), Gibco Earle’s balanced salt solution (EBSS) with calcium and magnesium (11540616), anti-LAMP1 (555798), anti-mouse Alexa Fluor^tm^ 633 (A21050), anti-Rabbit Alexa Fluor^tm^ 633 (A21070) and BCA kit (23227) were purchased from Thermo Fisher (Waltham, MA). Lenti-X packaging Singel Shots VSV-G (631276) were provided by Takara Bio Europe (Saint-Germain-en-Laye, France). JetOPTIMUS DNA Transfection Reagent (101000025) was purchased from VWR (Radnor, Pennsylvania).

### Plasmids and primers

The plasmids and primers used in this study are summarized in Supplementary Table 1.

### Generation of stably expressing cell lines

HexaKO, HeLa WT and HexaKO-LC3B cell lines were kindly provided by Dr. Sharon Tooze (Francis Crick institute, London). HexaKO-LC3C, HexaKO-LC3C G126A, HexaKO-LC3B G120A, HexaKO-LC3B/C, HexaKO-LC3C/B and HexaKO-RFP-LC3B stable cell lines were generated using Lentiviral transduction. The cDNA of interest was amplified by PCR and subcloned into pLVX-Puro plasmid by restriction enzyme cloning. The primers used are listed in supplementary table 1. LentiX cells were transfected with the VSV-G lentiviral packaging system (Takara Bio Europe, Saint-Germain-en-Laye, France) and the pLVX-puro vector containing the sequence of interest. After 48 h, the culture medium was collected and filtered with a 0.45 μm filter. HexaKO cells were treated with the lentivirus containing medium. After 24 h cells were incubated with full medium (FBS %10 and penicillin and streptomycin). Transduced clones were selected by adding 2 μg/mL of puromycin to the medium.

### Cell culture

Cells were grown in high glucose DMEM, supplemented with 10% fetal bovine serum (FBS), penicillin and streptomycin in a humidified atmosphere containing 5% CO₂ at 37 °C. For the stably expressing cell lines, the medium was supplemented with 1 μg/mL of puromycin to maintain selection. The experiments were performed without puromycin in the medium. Specific experimental conditions are indicated in the figure legends.

### Proteinase K assay

Cells were seeded in 150-mm dishes and cultured in either DMEM or EBSS for 6 h with 100 nM bafilomycin A1. Following treatment, cells were scraped in homogenization buffer (20 mM HEPES, 220 mM mannitol, 70 mM sucrose, 1 mM EDTA, pH 7.5). Cells were subsequently homogenized by passing them 40 times through a cell homogenizer, and the resulting lysates were clarified by centrifugation at 1,500 x g for 10 minutes. For the proteinase K protection assay, lysates were divided into three equal-volume aliquots and either left untreated, treated with 10 μg/mL proteinase K alone, or treated with 10 μg/mL proteinase K in the presence of 0.2% Triton X-100. Samples were incubated on ice for 30 min. Proteinase K activity was subsequently inhibited by boiling the samples for 10 min.

### Cell lysis

Cells were cultured in DMEM supplemented with FBS and antibiotics or with EBSS in the absence or presence of 100 nM of Bafilomycin A1 for 2, 4, and 6 h. After incubation, cells were washed with PBS and scraped in 120 μL of RIPA buffer. To remove cell debris and membranes, the samples were centrifuged for 10 minutes at 12000 x g. The supernatant was collected and protein concentration was measured with the BCA assay following the manufactureŕs instructions.

### Western Blot

30 μg of total protein per sample were loaded into an SDS-PAGE gel and transferred to a PVDF membrane with a Mini TransBlot© Cell system (Bio-Rad) (75 min, 400mA). The membrane was blocked with 5% non-fat milk. Membranes were incubated overnight at 4 °C with a 1:1000 dilution of the indicated antibody, except anti-NDP52, which was used at a 1:500 dilution. Membranes were then incubated with HRP-conjugated anti-mouse or anti-rabbit secondary antibodies (1:5000) and incubated for 1h at room temperature. Protein bands on the membranes were subsequently detected using Luminata Forte Western HRP Substrate. The signal was recorded in an iBright CL 1500 imaging system (Thermo Fisher scientific). Band intensities were quantified with Fiji (Image J) software.

### Immunofluorescence (IF)

Cells were cultured in DMEM or EBSS for 2 h and 6 h in coverslips (Fisher, 01175330). Then, they were fixed with 2% paraformaldehyde and permeabilized with 0.01% Triton X-100 for 10 minutes. The coverslips were then washed three times with PBS and blocked with 5% BSA in PBS. In the next step, cells were incubated with the primary antibodies diluted in 1% BSA/PBS for 1h at room temperature. The antibody dilutions can be found in Supplementary Table 2. After 3x PBS washes the coverslips were incubated with 1:500 dilution of anti-mouse or anti-rabbit Alexa Fluor^tm^ 633 (incubation 1h at room temperature). Nuclei were stained with DAPI (5 min 1:10,000 dilution). After 3x PBS washes and a last wash in deionized water, the coverslip was mounted on microscope slides (VWR, 631-1558) using Mowiol for its visualization in a Leica TCS SP5 II fluorescence microscope. For each independent experiment, at least 10 images were obtained, with a 63x/1.20 W HCX PL APO CS CORR objective and a 2.5 electronic zoom. Each image consisted of at least 10 cells. The laser wavelengths used were 405 nm for DAPI, 633 nm for Alexa Fluor 633 and 488 nm for GFP and the emission was recorded between 415-465 nm for DAPI, 500-530 nm for GFP and 650-800 nm for Alexa 633.

### Colocalization of GFP-tagged LC3 proteins with HexaKO-RFP-LC3B

HexaKO-RFP-LC3B stable cells were seeded in an 8 well ibiTreat chambered coverslip (Ibidi, 80826) and transfected either with GFP-LC3B, GFP-LC3C, GFP-LC3C/B, or GFP-LC3B/C. Transfection was performed using jetOPTIMUS (Polyplus, 101000025) following the manufacturer’s instructions and using a 1:2 (DNA:jetOPTIMUS) ratio. Prior to fixation, cells were either maintained in complete medium or incubated in EBSS for 2 h to induce starvation. 24 h after transfection cells were stained with DAPI and fixed in 2% paraformaldehyde for its visualization in a Leica TCS SP5 II fluorescence microscope. The laser wavelengths used were 405 nm for DAPI, 488 nm for GFP and 543 nm for RFP and the emission was recorded between 415-465 nm for DAPI, 500-530 nm for GFP and 570-650 nm for RFP.

### Live cell imaging

For live-cell imaging experiments, cells were incubated in EBSS during 1h and imaged using the same confocal microscope and acquisition settings as those used for fixed cells, but with an incubation chamber maintaining the temperature at 37 °C. Images were acquired with a scan time of 30 s, and the resulting video was exported at 10 frames per second.

### Fluorescence microscopy image analysis

The areas of GFP-positive puncta and p62 bodies were quantified using Fiji software (Image J). First, a Gaussian blur filter (σ = 0.5) was applied to the images, followed by thresholding to exclude background signal and non-specific fluorescence. In both cases, the threshold was manually adjusted for both GFP puncta and p62 bodies. The average particle area and the total number of particles per image were recorded. To determine the number of GFP puncta per cell, the total number of GFP puncta in each image was normalized to the number of nuclei in the corresponding image. Nuclei were identified by applying the Huang thresholding algorithm in Fiji to the DAPI channel, which had been smoothed five times beforehand. Only particles with a diameter of at least 10 μm were considered nucleus.

Colocalization analysis was performed using the object-based colocalization method implemented in the JaCoP plugin in FIJI. Threshold for both red and green channels was manually adjusted to exclude background signal. Colocalization percentage was calculated as the number of GFP-puncta colocalizing with RFP-puncta or Alexa 633 signal divided by the total amount of GFP puncta.

### Statistics

Western blot data are presented as the mean ± standard deviation (SD) of at least three independent experiments. Where appropriate, statistical significance was assessed using Student’s t-test. For immunofluorescence, 3 independent experiments were performed with at least 10 images taken for each, unless otherwise stated. Data are presented as the mean ± SD. Where appropriate, statistical significance was determined using one-way analysis of variance (ANOVA).

## RESULTS

### 1. LC3C forms more numerous and larger autophagic vesicles than LC3B

To investigate the functional differences between LC3B and LC3C in autophagy, HeLa cells lacking all six members of the LC3/GABARAP family (HexaKO) were stably reconstituted with either GFP-LC3B or GFP-LC3C, generating the HexaKO-LC3B and HexaKO-LC3C cell lines, respectively. This experimental approach allowed the behavior of each protein to be studied independently, in the absence of the other ATG8 family members.

First, the intracellular distribution of GFP-LC3B and GFP-LC3C was analyzed by confocal fluorescence microscopy under basal conditions (fed) and after starvation-induced autophagy. For this purpose, HexaKO-LC3B and HexaKO-LC3C cells were incubated in Earle’s Balanced Salt Solution (EBSS) for 2 or 6 h (Figure 1A), and the number and average area of GFP-positive puncta were quantified (Figure 1B, C). Under starvation conditions, both GFP-LC3B and GFP-LC3C were found in punctate structures distributed throughout the cytoplasm (Figure 1A). However, marked differences were observed between the two proteins. HexaKO-LC3C cells exhibited a significantly higher number of puncta per cell than HexaKO-LC3B cells. Moreover, the average area of GFP-LC3C-positive puncta was significantly larger than that of GFP-LC3B-positive puncta (Figure 1B, C). As a control, stable HexaKO cell lines expressing the non-lipidable mutants GFP-LC3C G126A and GFP-LC3B G120A were also generated (HexaKO-LC3C G126A and HexaKO-LC3B G120A cells, respectively). These mutants contain a substitution of the C-terminal glycine residue required for conjugation to phosphatidylethanolamine (PE), preventing their lipidation and association with autophagic membranes. These cell lines were used to determine whether the formation of GFP-positive puncta observed with LC3B and LC3C was dependent on their lipidation. Confocal microscopy analysis showed that both GFP-LC3C G126A and GFP-LC3B G120A exhibited a predominantly diffuse cytoplasmic distribution, indicating that puncta formation by LC3B and LC3C largely depends on their lipidation (Supplementary Figure 1).

**Figure 1.**
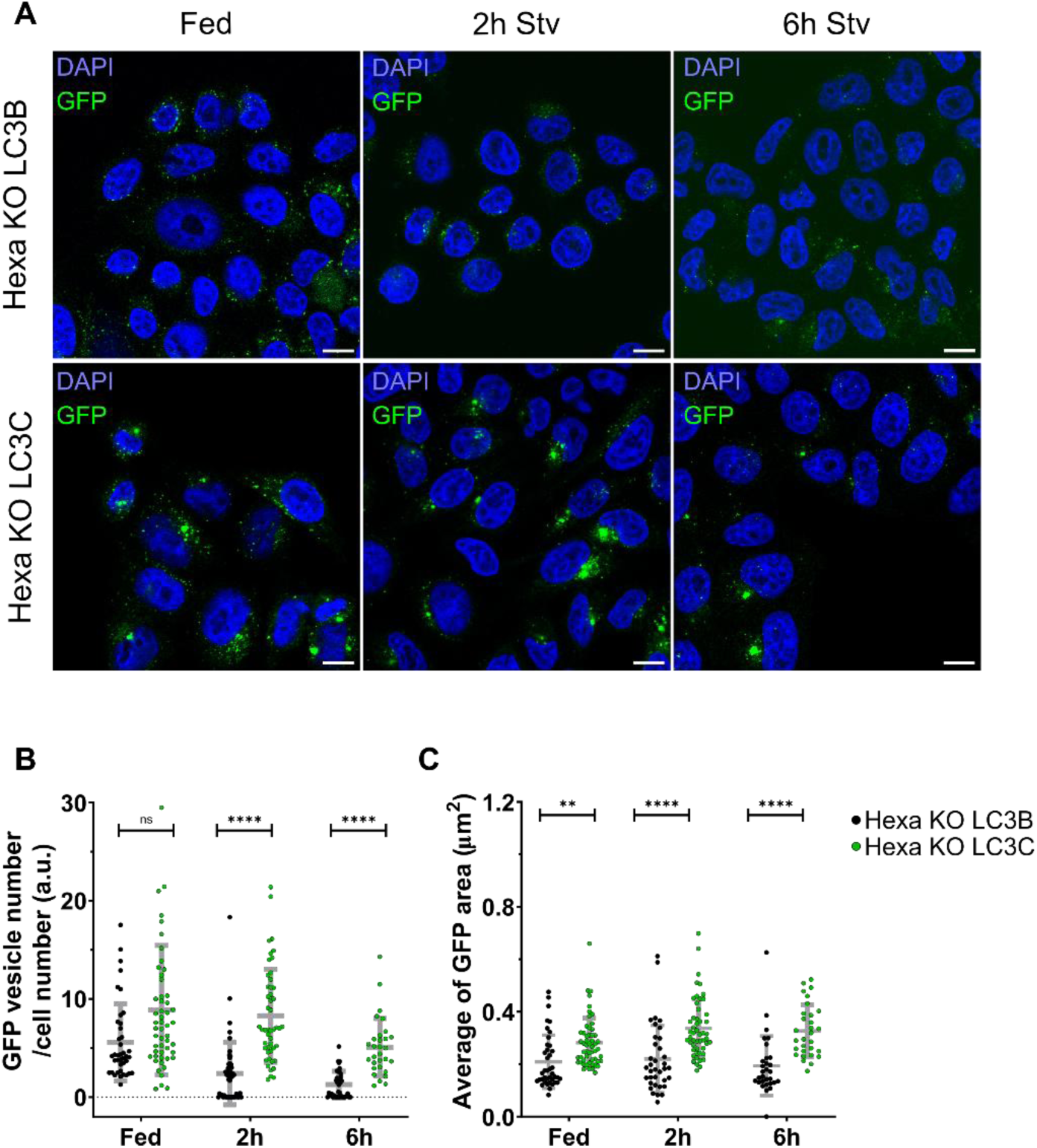
LC3C forms more numerous and larger puncta than LC3B. Representative fluorescence microscopy images of HexaKO-LC3B and HexaKO-LC3C in full media (fed) and after 2 h and 6 h of treatment with EBSS (A). Scale bar: 10 μm. Average number (B) and size (C) of GFP-labeled puncta. Average of at least 30 images ± SD. One-way ANOVA significance test: ****p<0.0001; ***p<0.001; ** p<0.01; * p<0.05; ns, non-significant.

These results indicate that LC3B and LC3C exhibit different behaviors during starvation-induced autophagy. LC3C appears to promote the formation or accumulation of more numerous and larger GFP-positive vesicles than LC3B. Furthermore, the absence of puncta in the non-lipidable mutants confirms that the formation of these vesicles depends on ATG8 lipidation.

### 2. LC3C forms functional autophagosomes under starvation conditions, although autophagosome closure is less efficient than with LC3B

#### 2.1. Both LC3B and LC3C undergo autophagic degradation in response to nutrient starvation

To further investigate the role of LC3B and LC3C in starvation-induced autophagy, the degradation of both proteins and autophagic flux were analyzed by western blotting. For this purpose, HexaKO-LC3B and HexaKO-LC3C cells were subjected to nutrient starvation, and GFP-LC3B and GFP-LC3C protein levels were analyzed at different time points. It should be noted that GFP-LC3C lipidation could not be reliably assessed by western blotting because the lipidated and non-lipidated forms were not sufficiently resolved by SDS-PAGE. The results showed that both LC3B and LC3C were degraded in response to starvation (Figure 2A, B, D, E). To determine whether this degradation was associated with autophagic flux, cells were treated with bafilomycin A1, an inhibitor of vacuolar H+-ATPase that prevents lysosomal acidification and autophagic degradation [31]. Autophagic flux of both proteins was assessed by comparing LC3B and LC3C levels in the presence and absence of bafilomycin A1. The results indicated that both LC3B and LC3C were degraded through lysosomal turnover, with significantly increased autophagic flux after 6 h of starvation compared with fed conditions (Figure 2C, F).

**Figure 2.**
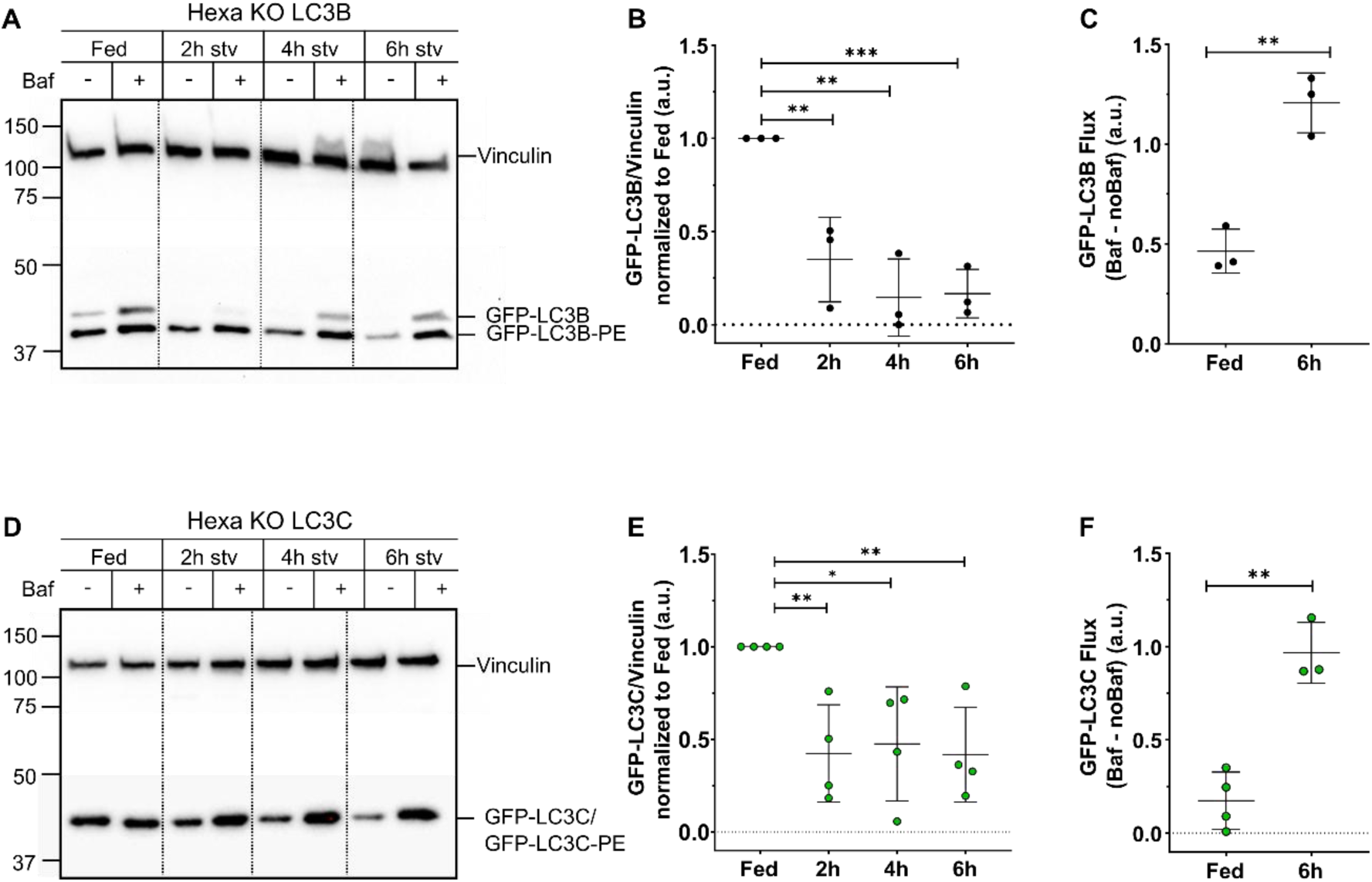
Both LC3B and LC3C undergo autophagic degradation in response to nutrient starvation. Representative western blots of HexaKO-LC3B (A) and HexaKO-LC3C (D) in full media (fed) and after 2 h, 4 h and 6 h of treatment with EBSS, with or without 100 nM of bafilomycin A1. Vinculin was used as a loading control. Quantification of GFP-LC3B (B) and GFP-LC3C (E) levels normalized to fed. GFP-LC3B (C) and GFP-LC3C (F) autophagic flux was calculated by subtracting GFP-LC3 levels in the absence of bafilomycin A1 from those measured in its presence. Values represent the mean ± SD of three independent experiments. Statistics were computed with Multiple Student’s t-test. ***p<0.001; ** p<0.01; * p<0.05; ns, non-significant.

The degradation of the non-lipidable mutants GFP-LC3B G120A and GFP-LC3C G126A was also analyzed in HexaKO-LC3B G120A and HexaKO-LC3C G126A cells under the same experimental conditions. Interestingly, both mutants showed a reduction in protein levels during starvation, but this degradation was not prevented by bafilomycin A1 treatment (Supplementary Figure 2A-F). These results suggest that the reduction in the levels of the non-lipidable mutants occurs independently of bafilomycin A1-sensitive lysosomal degradation and may involve autophagy-independent degradation pathways, such as the ubiquitin–proteasome system.

Both confocal microscopy and western blot analyses indicate that LC3C and LC3B, even in the absence of the other ATG8 family members, can participate in the formation of autophagic vesicles that undergo lysosomal degradation during starvation-induced autophagy. Furthermore, the different behavior of the non-lipidable mutants confirms the importance of ATG8 lipidation for their incorporation into autophagic membranes and subsequent autophagic degradation.

#### 2.2. Proteinase K protection assays indicate that LC3C allows autophagosome closure, although less efficiently than LC3B

To further investigate whether LC3C-positive autophagic structures could form functional autophagosomes in the absence of the other ATG8 family members, their ability to undergo autophagosome closure was assessed. For this purpose, proteinase K protection assays were performed in four cell lines: HeLa WT, HexaKO, HexaKO-LC3B and HexaKO-LC3C cells. Since autophagosome closure is impaired in HexaKO cells, whereas HeLa WT cells can form closed autophagosomes, these cell lines were used as negative and positive controls, respectively (Figure 3A, B). The proteinase K protection assay is based on the protection of autophagic cargo from proteolytic digestion when sequestered within closed autophagosomes. The addition of Triton X-100 was used as a positive control for proteinase K digestion, as membrane disruption exposes the protected cargo to the protease. p62 was selected as a marker of autophagic cargo, as previously described in similar assays [8]. To facilitate the detection and quantification of protected cargo, cells were treated with bafilomycin A1 to promote the accumulation of autophagic structures. The results showed that HexaKO-LC3C cells exhibited significantly higher protection of p62 against proteinase K digestion than HexaKO cells, indicating that LC3C is capable of promoting the closure of autophagosomes (Figure 3A–E). However, p62 protection was lower in HexaKO-LC3C cells than in HexaKO-LC3B cells. Interestingly, the degree of p62 protection observed in HexaKO-LC3B cells was comparable to that of HeLa WT cells, suggesting that LC3B alone can largely restore autophagosome closure in the absence of the other ATG8 family members. To complement these results, the protection of the ATG8 proteins themselves was also analyzed by detecting GFP-LC3B and GFP-LC3C by western blotting. Both proteins exhibited protection against proteinase K digestion, indicating that they could be incorporated into the inner membrane of closed autophagosomes (Figure 3C, D, F). To determine whether the ability of LC3B and LC3C to promote autophagosome closure depends on their lipidation, proteinase K protection assays were also performed using HexaKO-LC3B G120A and HexaKO-LC3C G126A cells, expressing the non-lipidable GFP-tagged mutants. Neither mutant exhibited protection of p62 or GFP against proteinase K digestion, indicating that the ability of LC3B and LC3C to promote autophagosome closure is dependent on their lipidation (Supplementary Figure 3). The proteinase K protection assays indicate that LC3C can participate in autophagosome closure, although less efficiently than LC3B. Nevertheless, LC3C retains sufficient capacity to promote the formation of closed autophagosomes, resulting in a significant improvement in autophagic function compared with cells lacking all ATG8 family members.

**Figure 3.**
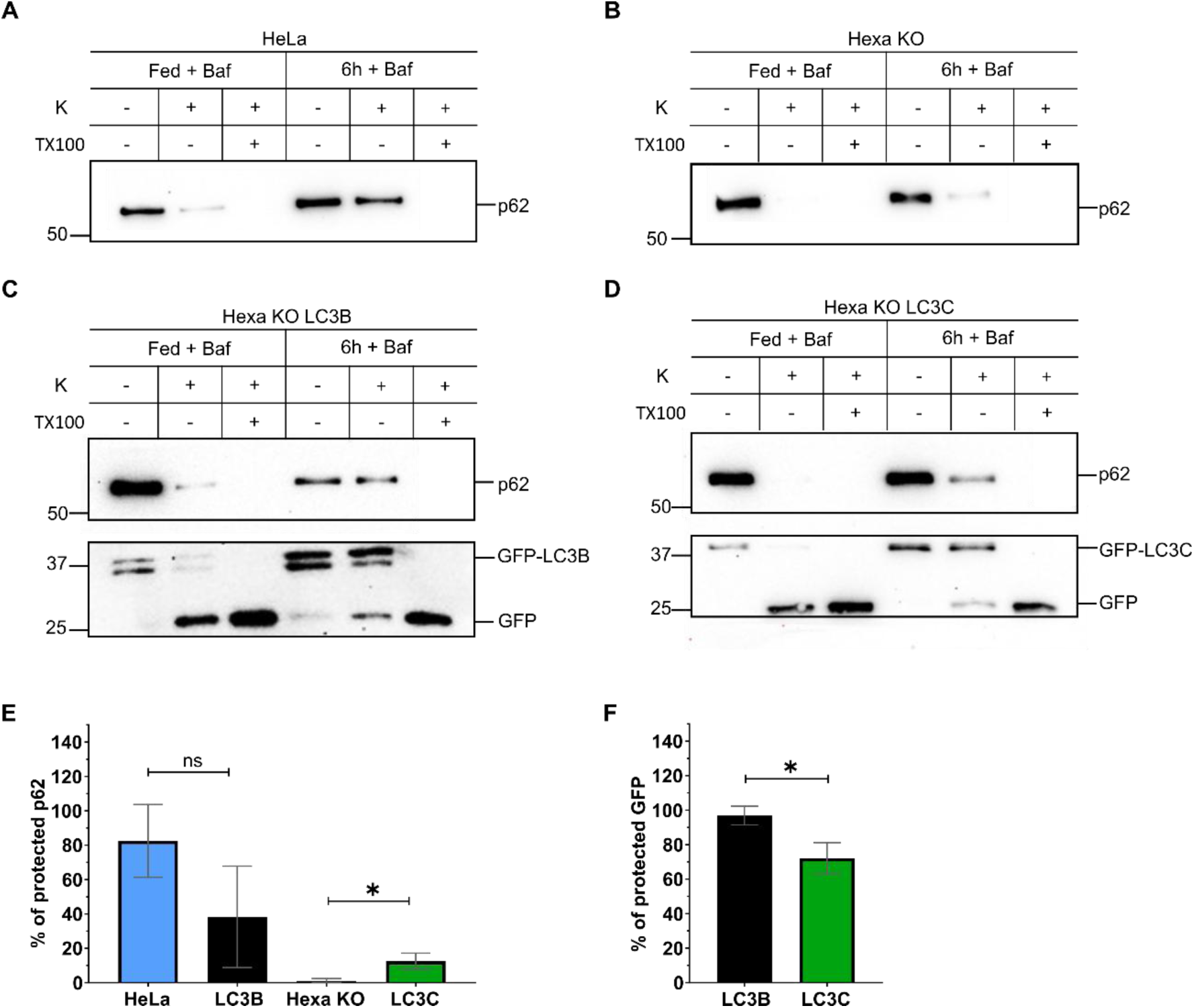
HexaKO-LC3B and HexaKO-LC3C can form closed autophagosomes. Representative Proteinase K assay western blots of HeLa (A), HexaKO (B), HexaKO-LC3B (C) and HexaKO-LC3C (D) in full media (fed) and after 6 h of treatment with EBSS and 100 nM of bafilomycin A1. Quantification of protected p62 (E) and protected GFP-LC3 (F) calculated using the following formula: (sample with proteinase K/ sample without proteinase K) x 100. Values represent the mean ± SD of three independent experiments. Statistical analysis was performed using Student’s t-test. ***p < 0.001; **p < 0.01; *p < 0.05; ns, not significant.

### 3. LC3B and LC3C differ in their ability to mediate the autophagic degradation of p62 and NDP52, suggesting receptor-specific functional specialization

#### 3.1. LC3B restores autophagic degradation of p62 in HexaKO cells, whereas LC3C shows a limited ability to rescue this phenotype

To further investigate the functional differences between LC3B and LC3C in the absence of the other ATG8 family members, the degradation of the autophagic receptor p62 during starvation-induced autophagy was analyzed. First, p62 degradation was compared in HeLa WT and HexaKO cells (Figure 4A, B). While p62 degradation was clearly observed in HeLa WT cells after 6 h of starvation, HexaKO cells exhibited defective p62 degradation (Figure 4A-C). Based on these results, HeLa WT and HexaKO cells were used as positive and negative controls, respectively, to determine whether LC3B or LC3C alone could restore p62 degradation. The results showed that stable expression of GFP-LC3B in HexaKO-LC3B cells partially restored p62 degradation after 6 h of starvation, although less efficiently than in HeLa WT cells (Figure 4C, D, F). This also confirmed that the N-terminal GFP tag did not prevent LC3B from mediating p62 degradation, validating the use of this experimental system to compare the functions of LC3B and LC3C. In contrast, GFP-LC3C in HexaKO-LC3C cells was unable to restore detectable p62 degradation under the same conditions, exhibiting a phenotype similar to that of HexaKO cells (Figure 4C, F). p62 degradation was also analyzed by western blotting in HexaKO-LC3B G120A and HexaKO-LC3C G126A cells stably expressing the corresponding non-lipidable GFP-tagged mutants. Neither mutant restored significant p62 degradation after 6 h of starvation, indicating that the ability of LC3B to mediate p62 degradation depends on its lipidation (Supplementary Figure 4).

**Figure 4.**
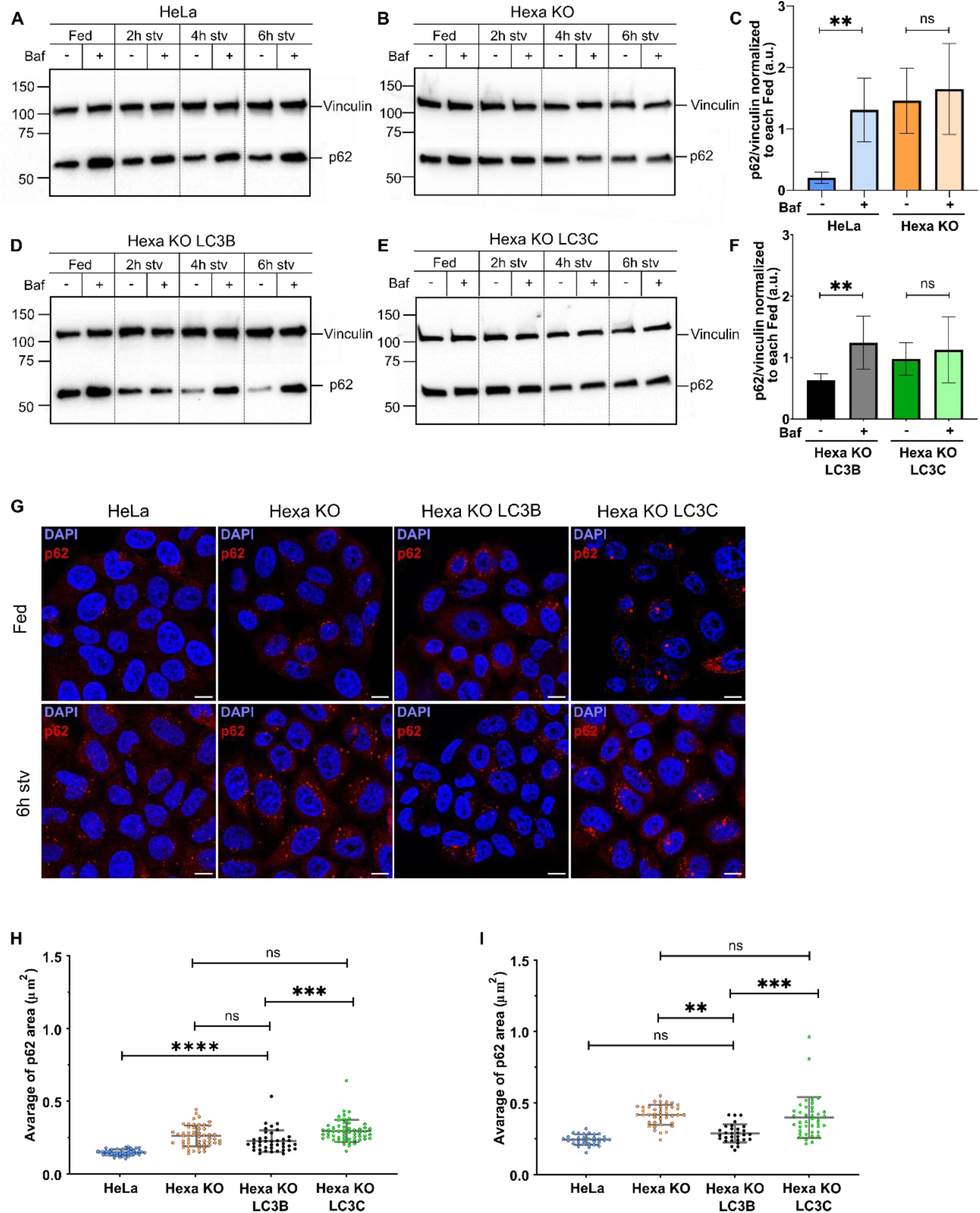
LC3B mediates p62 degradation more efficiently than LC3C. Representative western blots of HeLa (A), HexaKO (B), HexaKO-LC3B (D) and HexaKO-LC3C (E) in full media (fed) and after 2 h, 4 h, 6 h of treatment with EBSS with and without 100 nM of bafilomycin A1. Quantification of p62 levels after 6 h of starvation, normalized to Fed, in each cell line (C, F). Values represent the mean ± SD of at least three independent experiments. Statistics were computed with Student’s t-test. ***p<0.001; ** p<0.01; * p<0.05; ns, non-significant. (G) Representative fluorescence microscopy images of HeLa, HexaKO, HexaKO-LC3B and HexaKO-LC3C in full media (fed) and after 6 h of treatment with EBSS. Scale bar: 10 μm. Average size of p62 puncta in fed (H) and 6 h of starvation (I). Average of at least 30 images ± SD. One-way Anova significance test: ****p<0.0001; ***p<0.001; ** p<0.01; * p<0.05; ns, non-significant.

>To complement these results, the intracellular distribution of p62 was analyzed by immunofluorescence microscopy. p62 can accumulate in intracellular structures commonly referred to as p62 bodies, whose size can increase when autophagic degradation is impaired. Therefore, the area of p62-positive puncta was quantified in the four cell lines under basal conditions and after 6 h of starvation. Under fed conditions, HexaKO cells exhibited significantly larger p62 bodies than HeLa WT cells (Figure 4G, H), and after 6 h of starvation, the difference in p62 body size between HeLa WT and HexaKO cells was maintained (Figure 4G, I). Interestingly, HexaKO-LC3B cells were able to reduce p62 body size to levels comparable to those observed in HeLa WT cells after 6 h of EBSS treatment (Figure 4G, I). In contrast, HexaKO-LC3C cells exhibited p62 bodies of similar size to those found in HexaKO cells, indicating that LC3C was unable to rescue this phenotype (Figure 4G, I). Taken together, these results indicate that LC3B and LC3C differ in their ability to mediate p62 degradation in the absence of the other ATG8 family members. While LC3B partially restores p62 degradation and reduces the accumulation of large p62 bodies during starvation, LC3C fails to recover these functions efficiently. Thus, HexaKO-LC3B cells exhibit a phenotype closer to HeLa WT cells, whereas HexaKO-LC3C cells resemble HexaKO cells, suggesting a specific limitation of LC3C in the autophagic degradation of p62.

#### 3.2. LC3C mediates NDP52 degradation more efficiently than LC3B under starvation conditions

The results obtained with p62 raised the question of whether LC3C alone is capable of mediating selective autophagic degradation of receptors or whether the presence of other ATG8 family members is required for this function. To address this question, the degradation of NDP52, an autophagic receptor previously reported to interact preferentially with LC3C during xenophagy [14], was analyzed under the same starvation conditions used in the previous experiments. The results showed that NDP52 was significantly degraded in HeLa WT cells after 6 h of starvation, whereas its autophagic degradation was impaired in HexaKO cells (Figure 5A, B, E). As described above for p62, rescue experiments with GFP-LC3B or GFP-LC3C were performed to investigate whether NDP52 degradation was restored in HexaKO-LC3B and HexaKO-LC3C cells. Interestingly, GFP-LC3C was able to restore NDP52 degradation under starvation conditions, whereas GFP-LC3B failed to rescue this phenotype (Figure 5C–E). Although a reduction in NDP52 levels was also observed in HexaKO and HexaKO-LC3B cells during starvation, this reduction was not prevented by bafilomycin A1 treatment. These results suggest that NDP52 may be degraded through alternative, autophagy-independent pathways, such as proteasomal degradation.

**Figure 5.**
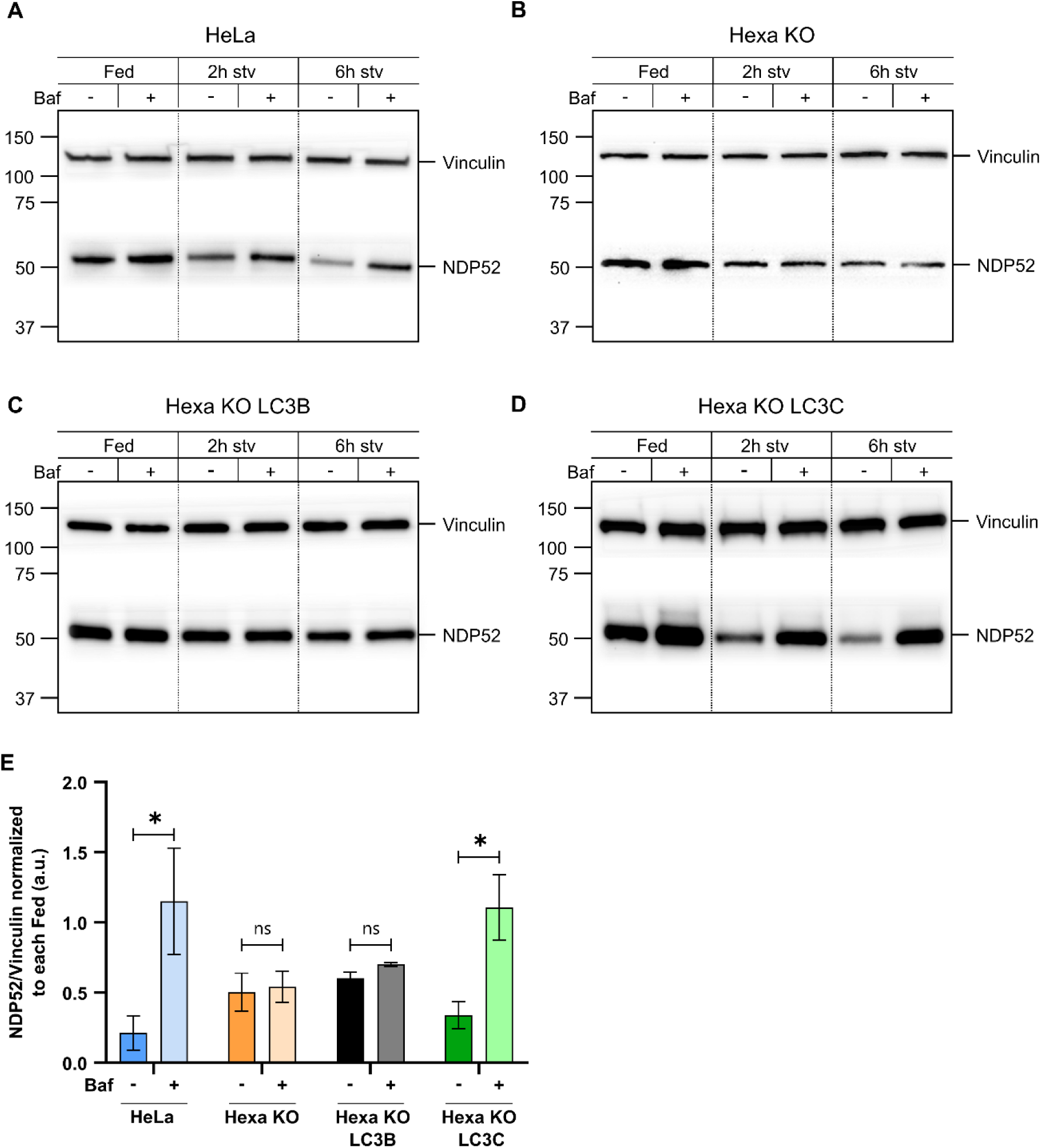
LC3C mediates NDP52 degradation more efficiently than LC3B under starvation conditions. Representative western blots of HeLa (A), HexaKO (B), HexaKO-LC3B (C) and HexaKO-LC3C (D) in full media (fed) and after 2 h and 6 h of treatment with EBSS with and without 100 nM of bafilomycin A1. Quantification of NDP52 levels after 6 h of starvation, normalized to Fed, in each cell line (E). Values represent the mean ± SD of at least three independent experiments. Statistics were computed with Student’s t-test. ***p<0.001; ** p<0.01; * p<0.05; ns, non-significant.

Overall, these results indicate that both LC3B and LC3C are able to mediate the autophagic degradation of specific receptors independently of the other ATG8 family members. However, their efficiency depends on the receptor involved. While LC3B restores p62 degradation but not NDP52 degradation, LC3C shows the opposite behavior, restoring NDP52 degradation but not p62 degradation. These findings suggest that LC3B and LC3C have developed specialized functions in selective autophagy, allowing them to mediate the degradation of different autophagic receptors without requiring the presence of other ATG8 proteins.

#### 4. LC3B and LC3C exhibit distinct colocalization patterns with components of the autophagic machinery

Considering the differences observed in the properties of LC3B- and LC3C-positive vesicles, as well as their distinct abilities to mediate the degradation of autophagic receptors, the intracellular colocalization patterns of these two proteins were investigated. For this purpose, the colocalization of GFP-LC3B and GFP-LC3C with two proteins associated with different stages of the autophagic process was analyzed by immunofluorescence: ATG9A, a transmembrane protein involved in the early stages of autophagosome formation, and LAMP1, a lysosomal marker associated with the final stages of autophagy. First, the colocalization of GFP-LC3B and GFP-LC3C with ATG9A was analyzed by confocal microscopy, under both fed and starvation conditions. The results showed that LC3C exhibited significantly higher colocalization with ATG9A than LC3B under both conditions (Figure 6A, B). The colocalization of both proteins with LAMP1 was then analyzed to investigate their association with lysosomal compartments or autophagolysosomes. In contrast to the results obtained with ATG9A, GFP-LC3B and GFP-LC3C showed similar levels of colocalization with LAMP1, with no significant differences between them (Figure 6C, D). These results indicate that LC3B and LC3C exhibit different colocalization patterns with components of the autophagic machinery. The higher colocalization of LC3C with ATG9A suggests that LC3C may have a distinct role in the early stages of autophagosome formation. In contrast, the similar levels of colocalization with LAMP1 suggest that both proteins can reach lysosomal compartments despite their different association with ATG9A.

**Figure 6.**
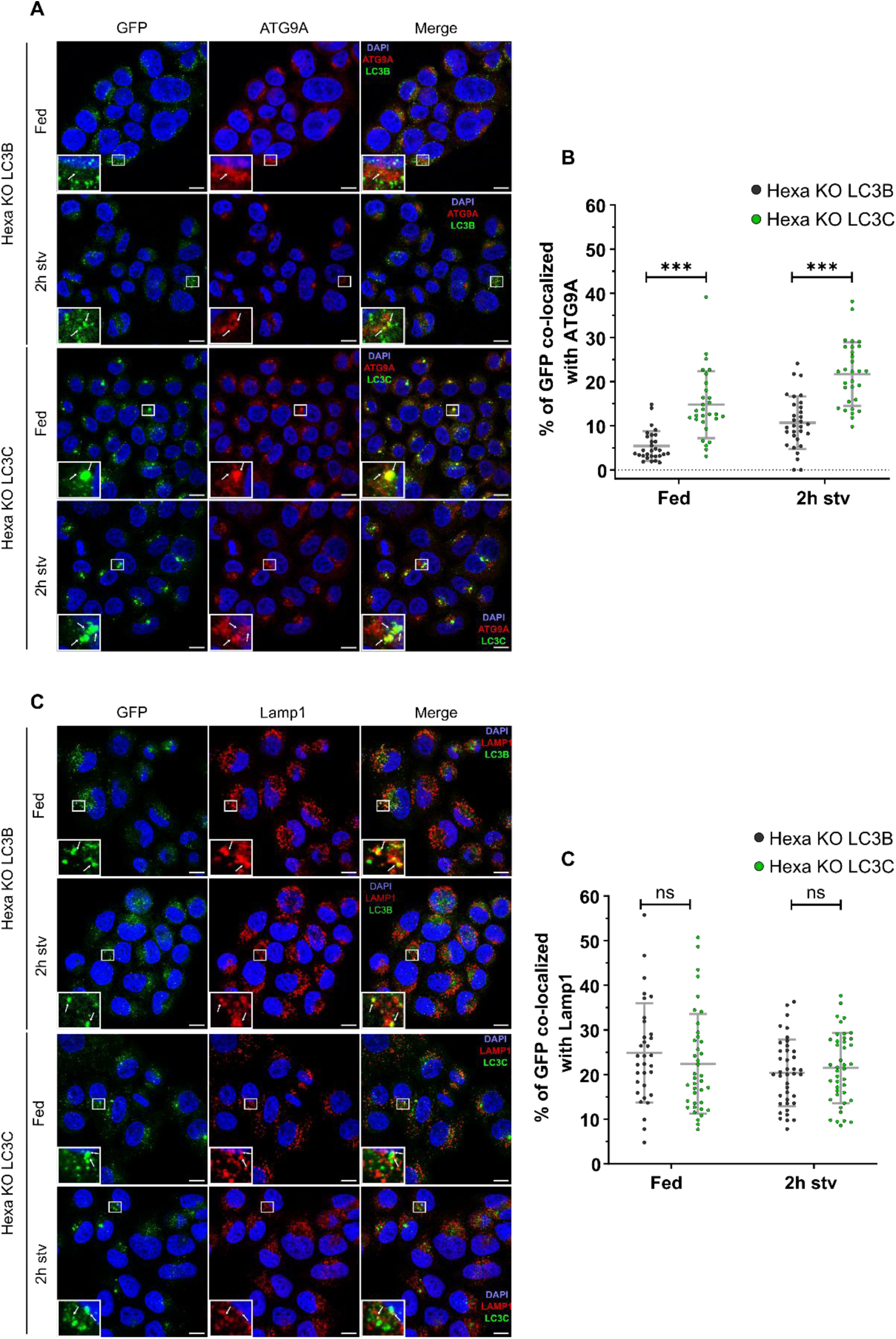
LC3B and LC3C show distinct colocalization patterns with autophagy-related proteins. Representative fluorescence microscopy images of HexaKO-LC3B and HexaKO-LC3C in full media (fed) and after 2 h of treatment with EBSS stained for ATG9A (A) and LAMP1 (C). Scale bar: 10 μm. Colocalization was expressed as the percentage of GFP-positive puncta colocalizing with each of the indicated markers. Colocalization percentage of GFP-LC3 with ATG9A (B) and GFP-LC3 with LAMP1 (D). Average of at least 30 images ± SD. One-way ANOVA significance test: ****p<0.0001; ***p<0.001; ** p<0.01; * p<0.05; ns, non-significant.

#### 5. LC3B and LC3C show limited colocalization, suggesting the existence of distinct autophagosome subpopulations

The differences observed between GFP-LC3B and GFP-LC3C in the size and number of their vesicles, their ability to mediate the degradation of specific autophagy receptors, and their colocalization with components of the autophagic machinery raised the question of whether these two proteins are incorporated into the same autophagosomes. To investigate this possibility, the colocalization of LC3B and LC3C was analyzed in cells expressing both proteins simultaneously. For this purpose, a stable HexaKO-RFP-LC3B cell line was generated, which was subsequently transiently transfected with either GFP-LC3B or GFP-LC3C (Figure 7A). The combination of GFP-LC3B and RFP-LC3B was used as a positive control, since both fluorescent proteins were fused to the same protein and were therefore expected to show a high degree of colocalization. Confocal microscopy analysis showed that, as expected, GFP-LC3B and RFP-LC3B exhibited a high degree of colocalization under both fed and starvation conditions (Figure 7B, C). In contrast, GFP-LC3C showed markedly lower colocalization with RFP-LC3B under both conditions, with significant differences compared with the GFP-LC3B/RFP-LC3B positive control (Figure 7B, C). Despite the presence of some vesicles positive for both proteins, colocalization analysis indicates that LC3B and LC3C are predominantly associated with different intracellular vesicles, even when both proteins are expressed in the same cells. To further investigate this observation, live-cell imaging experiments were performed to analyze the dynamics of LC3B- and LC3C-positive vesicles. These experiments confirmed the limited colocalization observed by confocal microscopy, as GFP-LC3C and RFP-LC3B-positive vesicles showed largely independent distributions and dynamics, with very few colocalization events detected over time (Supplementary Video 1). Our findings suggest that, in the absence of other ATG8 proteins, LC3B and LC3C are preferentially incorporated into distinct populations of autophagosomes, rather than being equally distributed among the same vesicles, which may contribute to their functional specialization in the autophagic process.

**Figure 7.**
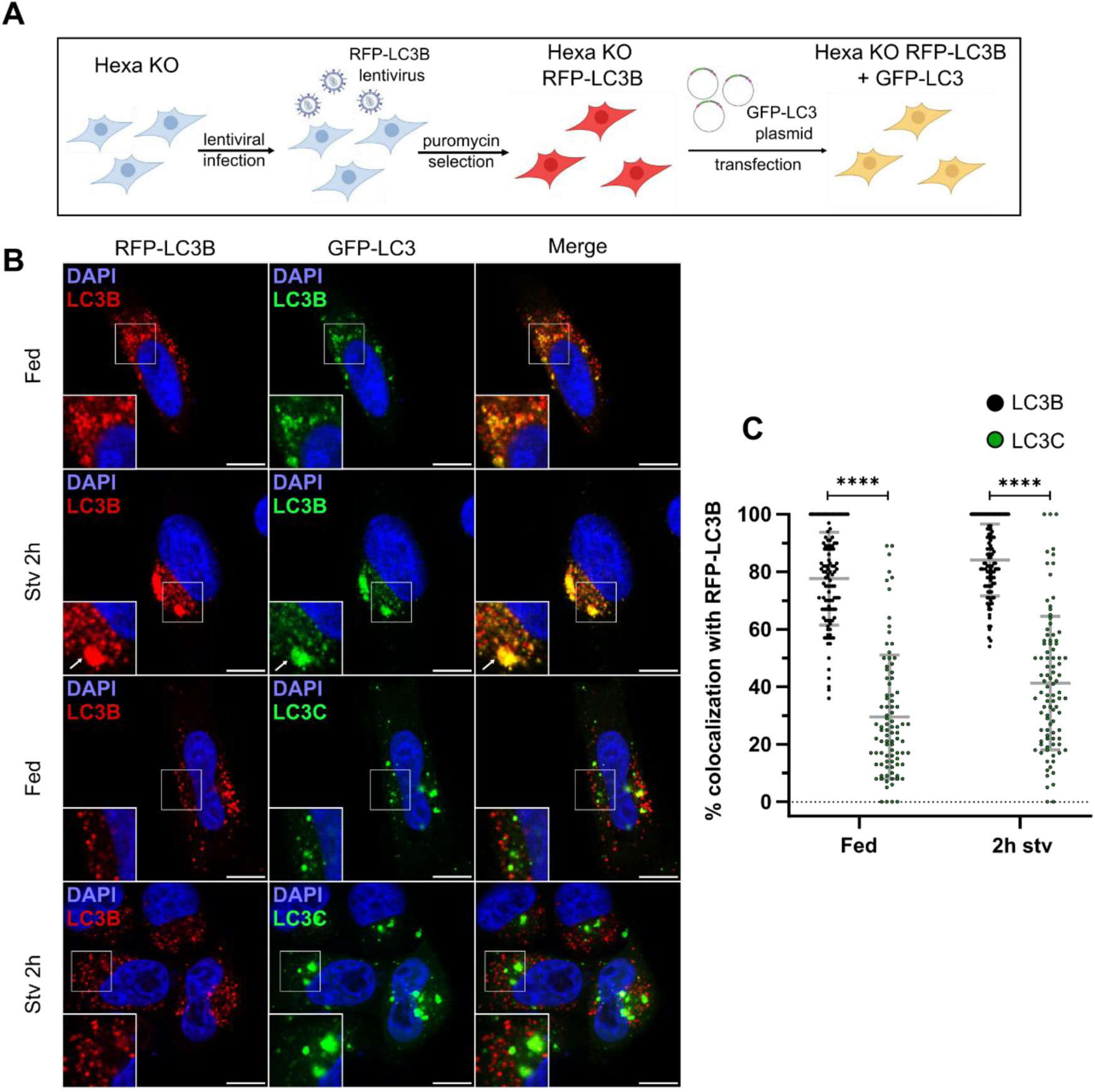
LC3B and LC3C localize to distinct populations of intracellular vesicles. Schematic representation of the experimental protocol (A). Representative fluorescence microscopy images of HexaKO-RFP-LC3B cells transfected either with GFP-LC3B or GFP-LC3C, in full media (fed) and after 2 h of treatment with EBSS (B) Scale bar: 10 μm. Colocalization percentage between LC3B and LC3C (C). Average of at least 30 images ± SD. One-way ANOVA significance test: ****p<0.0001; ***p<0.001; ** p<0.01; * p<0.05; ns, non-significant.

### 6. The N-terminal region of LC3C determines autophagic vesicle size, while its ubiquitin-like core affects intracellular localization

#### 6.1. The N-terminal region of LC3C determines autophagic vesicle size

In a previous study from our group, the membrane fusion properties of LC3B and LC3C were investigated using purified recombinant proteins and model membranes. These experiments showed that LC3C exhibited higher membrane fusion activity than LC3B and identified its N-terminal region as responsible for this difference [30]. The N-terminal region of ATG8 family proteins is conserved and generally composed of two α-helices. However, in LC3C, the region corresponding to the first N-terminal α-helix of LC3B adopts a disordered conformation. To investigate the contribution of this region to membrane fusion, two chimeric proteins were previously generated by exchanging the N-terminal regions of LC3B and LC3C. The first chimera, LC3C/B, contained the N-terminal region of LC3C followed by the ubiquitin-like core of LC3B. Conversely, the second chimera, LC3B/C, contained the N-terminal region of LC3B combined with the ubiquitin-like core of LC3C (Figure 8A). This experimental approach allowed the contribution of the N-terminal region and the ubiquitin-like core of each protein to their membrane fusion properties to be investigated. In the present study, LC3C was observed to form larger vesicles than LC3B in cells (Figure 1). Considering our previous findings, it was hypothesized that the increased size of LC3C-positive vesicles could be related to the higher membrane fusion activity of LC3C observed *in vitro*. To investigate this possibility, stable HexaKO cell lines expressing GFP-tagged versions of the same LC3C/B and LC3B/C chimeras used in our previous study were generated (HexaKO-LC3C/B and HexaKO-LC3B/C cells, respectively; Figure 8A). The size of the GFP-positive puncta formed by these proteins was then analyzed by confocal microscopy, under both fed and starvation conditions. The results showed that GFP-LC3C/B formed puncta of similar size to those observed in HexaKO-LC3C cells, whereas GFP-LC3B/C formed puncta of similar size to those observed in HexaKO-LC3B cells. This behavior was maintained under both fed and starvation conditions (Figure 8B, C). These findings indicate that the N-terminal region of LC3C is a key determinant of the increased size of LC3C-positive vesicles in cells. Moreover, the agreement between the results obtained with model membranes in our previous study and those observed in cells suggests that the membrane fusion properties of LC3C may contribute to the formation of larger autophagic vesicles.

**Figure 8.**
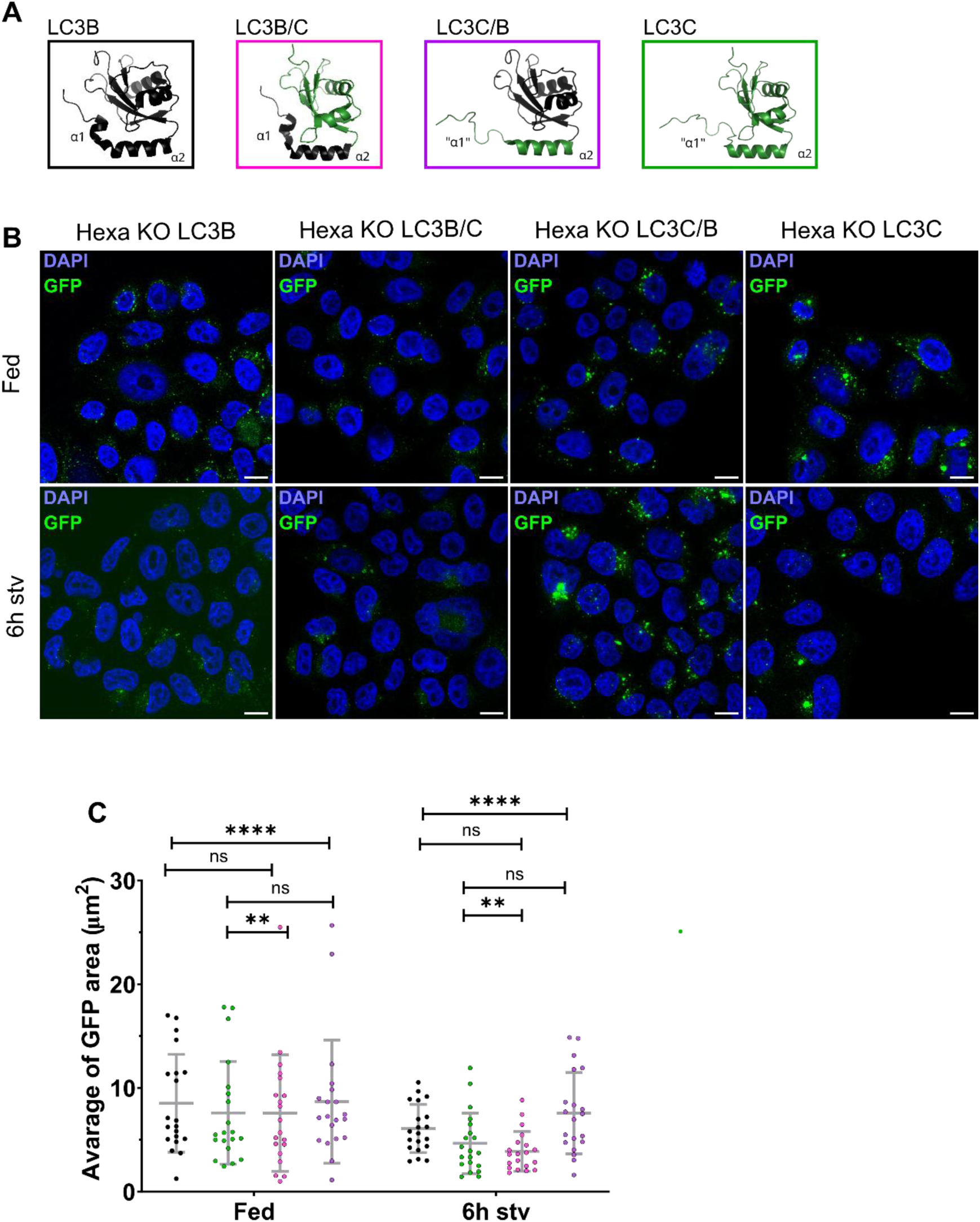
The N-terminal region of LC3C determines the size of GFP-positive puncta. LC3B (2ZJD) and LC3C (2NCN) structures displayed by PyMOL and the proposed structures of the chimeras LC3B/C (N-terminus of LC3B and ubiquitin-like core of LC3C) and LC3C/B (N-terminus of LC3C and ubiquitin-like core of LC3B) (A). Representative fluorescence microscopy images of HexaKO-LC3B, HexaKO-LC3B/C, HexaKO-LC3C/B and HexaKO-LC3C in full media (fed) and after 6 h of treatment with EBSS (B). Scale bar: 10 μm. Average size (C) of GFP-labeled puncta. Average of at least 20 images ± SD. One-way ANOVA significance test: ****p<0.0001; ***p<0.001; ** p<0.01; * p<0.05; ns, non-significant.

To further investigate the functional properties of the chimeric proteins, their ability to reduce the size of p62 bodies during starvation-induced autophagy was analyzed. As described above, GFP-LC3B was more efficient than GFP-LC3C in reducing the size of p62 bodies, restoring a phenotype similar to that observed in HeLa WT cells (Figure 4). To determine whether this difference was associated with the N-terminal region or the ubiquitin-like core, the size of p62-positive puncta in HexaKO-LC3C/B and HexaKO-LC3B/C cells was analyzed by confocal microscopy. The results showed that neither chimera was able to reduce the size of p62 bodies to the levels observed in HexaKO-LC3B cells (Supplementary Figure 5B, C). These findings suggest that both the N-terminal region and the ubiquitin-like core of LC3B are required for its ability to efficiently reduce the accumulation of large p62 bodies during starvation.

#### 6.2. The ubiquitin-like core determines the distinct colocalization patterns of LC3B and LC3C

The possible contribution of the N-terminal region of LC3C to the differences in intracellular localization observed between LC3B and LC3C was next investigated. For this purpose, the stable HexaKO-RFP-LC3B cell line described above was transiently transfected with GFP-tagged versions of the LC3C/B and LC3B/C chimeras. The colocalization of both chimeric proteins with RFP-LC3B was then analyzed by confocal microscopy (Figure 9A–C). Interestingly, the results showed that, in contrast to the differences observed in vesicle size, the colocalization patterns of the chimeric proteins were determined by their ubiquitin-like cores. GFP-LC3C/B, containing the N-terminal region of LC3C and the ubiquitin-like core of LC3B, exhibited high colocalization with RFP-LC3B, similar to that observed with GFP-LC3B. In contrast, GFP-LC3B/C, containing the N-terminal region of LC3B and the ubiquitin-like core of LC3C, showed limited colocalization with RFP-LC3B, resembling the behavior of GFP-LC3C (Figure 9A-C). These results indicate that the N-terminal region and the ubiquitin-like core of LC3B and LC3C play distinct roles in determining their cellular behavior. While the N-terminal region of LC3C is responsible for the increased size of LC3C-positive vesicles, the ubiquitin-like core appears to determine the distinct colocalization patterns of LC3B and LC3C. In contrast, neither chimeric protein was able to reproduce the ability of LC3B to reduce the size of p62 bodies, suggesting that both regions of LC3B are required for this function. These findings suggest that the membrane-remodeling properties of LC3C observed *in vitro* may contribute to the formation of larger autophagic vesicles in cells, while other functions, such as p62 body clearance and intracellular localization, depend on different structural determinants.

**Figure 9.**
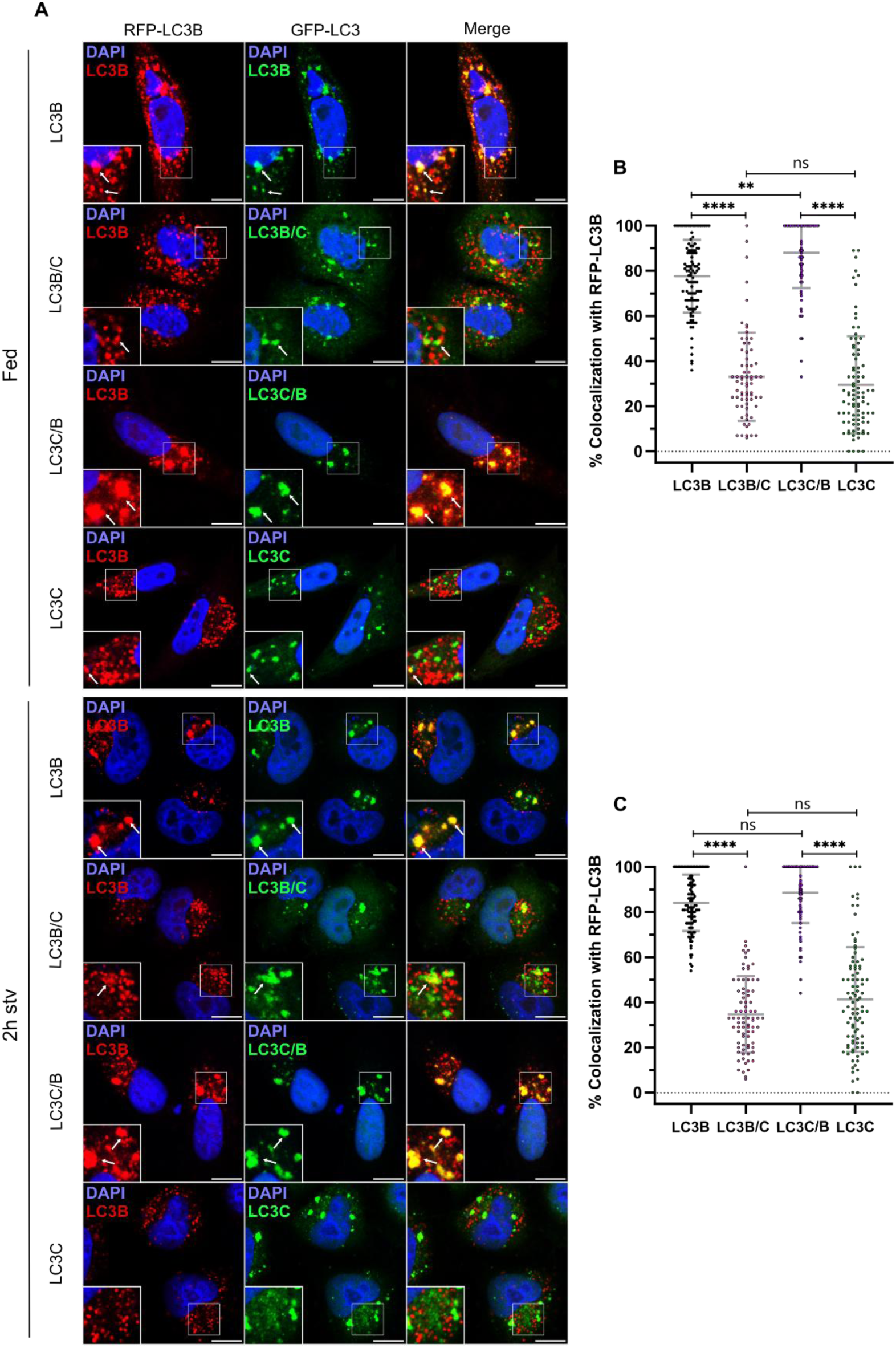
The ubiquitin-like core determines the distinct colocalization patterns of LC3B and LC3C. Representative fluorescence microscopy images of HexaKO-RFP-LC3B cells transfected either with GFP-LC3B, GFP-LC3B/C, GFP-LC3C/B or GFP-LC3C, in full media (fed) and after 2 h of treatment with EBSS (A) Scale bar: 10 μm. Colocalization percentage between RFP-LC3B with GFP-LC3B, GFP-LC3B/C, GFP-LC3C/B or GFP-LC3C in fed (B) and after 2 h of starvation (C). Average of at least 30 images ± SD. One-way ANOVA significance test: ****p<0.0001; ***p<0.001; ** p<0.01; * p<0.05; ns, non-significant.

## DISCUSSION

In this study, we investigated the functional specialization of LC3C within the ATG8 protein family. For this purpose, we compared the autophagic properties of LC3C and LC3B using HeLa WT cells and ATG8 HexaKO cells individually reconstituted with either protein. Our results reveal important differences between LC3B and LC3C in autophagic vesicle formation, cargo degradation, and intracellular localization, indicating that these proteins have developed distinct functional properties.

Several aspects of our experimental system should be taken into account when interpreting the results. Although ATG8-positive puncta are widely used as autophagosome markers [32], ATG8 proteins also participate in non-canonical processes involving single-membrane compartments [33]. Thus, not all GFP-positive puncta necessarily represent autophagosomes. Nevertheless, their reduction during starvation, together with the bafilomycin A1-sensitive degradation of GFP-LC3B and GFP-LC3C (Figure 2), indicates that a substantial fraction is associated with canonical autophagy. Puncta size remained relatively constant under fed and starvation conditions despite reduced puncta numbers and protein levels after 6 h of starvation (Figure 1), suggesting that the size differences between LC3B and LC3C do not simply reflect protein abundance. Although GFP is a relatively large tag, both fusion proteins retained autophagic activity, as shown by their contribution to autophagosome closure and receptor degradation in LC3/GABARAP HexaKO cells (Figure 3, 4, 5). These results indicate that both GFP-tagged proteins retain their functionality under our experimental conditions.

Overexpression of GFP-tagged ATG8 proteins can generate aggregates that resemble autophagic puncta [34]. However, non-lipidable LC3B G120A and LC3C G126A mutants showed a predominantly diffuse cytoplasmic distribution (Supplementary Figure 1), indicating that puncta formation largely depends on lipidation. Neither mutant restored autophagosome closure or receptor degradation in LC3/GABARAP HexaKO cells, further indicating that the functions observed require ATG8 membrane conjugation.

Although six ATG8 proteins are generally recognized as the major members of the human ATG8 family, two additional genes, MAP1LC3B2 and GABARAPL3, have also been described [11]. GABARAPL3 is classified as a pseudogene, with no demonstrated contribution to autophagy, whereas MAP1LC3B2 encodes a protein highly similar to LC3B. However, the physiological relevance of LC3B2 remains poorly understood, and its contribution to endogenous autophagic activity has not been established [35]. Importantly, although MAP1LC3B2 might not have been specifically targeted during the generation of the ATG8 HexaKO cells, no LC3B antibody signal was detected in these cells. This observation is consistent with low or undetectable LC3B2 protein levels. Furthermore, the pronounced defects in autophagic degradation observed in HexaKO cells indicate that any potential endogenous LC3B2 expression is insufficient to compensate for the absence of the six major ATG8 proteins. Thus, the HexaKO model provides a suitable experimental system to investigate the individual contributions of LC3B and LC3C to autophagosome formation and function, without substantial functional compensation by other canonical ATG8 family members.

Compared to the wild-type HeLa cells, LC3/GABARAP HexaKO cells showed defective autophagosome closure, impaired starvation-induced degradation of p62 and NDP52, and accumulation of p62 bodies (Figure3, 4, 5). Reconstitution with either LC3B or LC3C partially restored autophagic function, but with distinct efficiencies: LC3B promoted closure and p62 degradation more effectively, whereas LC3C was more effective in restoring NDP52 degradation. Thus, individual ATG8 proteins can mediate distinct autophagic functions without the other family members. Apart from the differences in their ability to restore autophagy, LC3C and LC3B showed limited colocalization with each other and distinct colocalization patterns with other autophagy markers, suggesting that they may be incorporated into different autophagosome populations.

Both LC3B and LC3C sustained starvation-induced autophagic flux when expressed individually in LC3/GABARAP HexaKO cells, as indicated by bafilomycin A1-sensitive protein degradation, reduced GFP-positive puncta during starvation, and colocalization with LAMP1 (Figure 1 and 6). This is particularly relevant, given the prominent role attributed to GABARAP proteins in starvation-induced autophagy [8,9]. Although GABARAPs may contribute more strongly to autophagic efficiency, our findings indicate that LC3B and LC3C retain common basic functions that allow them to sustain functional autophagy independently of other ATG8 family members, despite the evolutionary specialization of the ATG8 family [36].

Autophagosome closure involves sealing of the phagophore membrane, a process mediated primarily by ESCRT machinery, including ESCRT-III and VPS4 [37,38]. ATG8 proteins may also maintain autophagosomal membrane integrity through interactions with ESCRT components [39]. In our system, either LC3B or LC3C partially restored the proteinase K protection lost in LC3/GABARAP HexaKO cells, although LC3C was less efficient (Figure 3). The reason for this difference remains unclear, but it could reflect differences in the ability of LC3B and LC3C to recruit or interact with components of the autophagosome closure machinery. Differences in p62 sequestration into LC3B- and LC3C-positive autophagosomes may also contribute to the different levels of p62 protection observed in these cells. Overall, these findings suggest that LC3B and LC3C differ in their efficiency in promoting autophagosome closure, consistent with functional specialization within the ATG8 family.

LC3B and LC3C also differed in their ability to mediate receptor degradation. Although starvation-induced degradation of both p62 and NDP52 was impaired in ATG8 Hexa-KO cells, LC3B restored p62 degradation more efficiently, whereas LC3C preferentially restored NDP52 degradation (Figure 4, 5). This receptor-dependent behaviour indicates functional specialization even in the absence of other ATG8 family members. p62 is a major receptor for the selective autophagic removal of ubiquitinated protein aggregates and contributes to cellular proteostasis [40,41]. LC3/GABARAP Hexa-KO cells accumulated large p62-positive bodies, and LC3C restored their degradation only to a limited extent. This agrees with Nguyen et al. [8], but contrasts with Liu et al. [16], who implicated LC3C in aggrephagy through its interaction with WDR81. Unlike our system, those experiments were performed in cells containing other ATG8 proteins. LC3C may therefore require cooperation with other family members for efficient p62 degradation, although the underlying mechanism remains unknown. NDP52, encoded by CALCOCO2, mediates selective autophagy of intracellular pathogens and damaged mitochondria through xenophagy and mitophagy [42,43]. In the work by von Muhlinen et al. [14] a non-canonical LC3-interacting region (CLIR) that preferentially binds LC3C was identified in NDP52. Consistent with this interaction, LC3C restored starvation-induced NDP52 degradation more efficiently than LC3B in ATG8 Hexa-KO cells (Figure 5). Our findings extend this preferential relationship beyond the selective pathways in which it was initially described and indicate that LC3C can mediate NDP52 turnover independently of other ATG8 proteins.

ATG9A is a transmembrane protein involved in autophagosome biogenesis. ATG9A-containing vesicles contribute to the initial membrane from which autophagosomes develop [44]. Furthermore, its lipid scramblase activity contributes to autophagosomal membrane expansion [45,46], and ATG9A is also necessary for proper autophagic function [47]. In our experiments, LC3C-positive puncta showed greater colocalization with ATG9A than those formed by LC3B, both under fed and starved conditions. This difference may suggest that LC3C is more frequently associated with ATG9A-containing compartments, potentially reflecting differences in the intracellular distribution or maturation of LC3B- and LC3C-positive autophagic vesicles. However, colocalization analysis alone does not establish a direct interaction or a specific functional relationship between LC3C and ATG9A, and further experiments would be required to clarify the significance of this observation.

An important finding of this study is the limited colocalization between LC3B- and LC3C-positive puncta, suggesting that these proteins preferentially associate with different populations of autophagic vesicles. This observation was confirmed by live-cell imaging experiments (Supplementary Video 1), which revealed that LC3B- and LC3C-positive puncta remained largely separated during vesicle movement, with only occasional colocalization events. Importantly, the high degree of colocalization observed between GFP-LC3B and RFP-LC3B in control experiments indicates that the limited overlap between LC3B and LC3C is not simply a consequence of the different sensitivities of the two fluorophores to acidic environments. Although several studies have described functional differences among ATG8 family members [11,48], comparatively few studies have examined whether this functional diversity is reflected in the existence of distinct autophagosome populations. Interestingly, Goldin-Azulay et al. [49] recently demonstrated that different members of the GABARAP subfamily preferentially associate with distinct populations of autophagic vesicles. Furthermore, their proteomic analysis revealed differences in the protein composition of vesicles associated with individual GABARAP members, suggesting that these populations may have distinct cellular functions. Our results extend these observations to LC3B and LC3C, suggesting that the differential distribution of ATG8 family members may contribute to the existence of distinct autophagosome populations. Although the functional properties of these populations remain to be established, their differential association with LC3B or LC3C could represent an additional level of functional specialization within the ATG8 family. However, it is important to consider that these experiments were performed in LC3/GABARAP HexaKO cells expressing only LC3B and LC3C. Therefore, we cannot exclude the possibility that the absence of other ATG8 family members may influence the distribution of these proteins and accentuate the differences observed between LC3B- and LC3C-positive vesicles. Further studies examining the localization of endogenous ATG8 proteins will be necessary to determine whether this differential distribution is maintained under physiological conditions.

One of the most interesting results of this study is that the N-terminal region of LC3C is a key determinant of the increased size of LC3C-positive autophagic vesicles compared with those formed by LC3B (Figure 8). ATG8 proteins contain two N-terminal α-helices, although LC3C exhibits distinctive structural features in this region, including six additional amino acids and a disordered segment corresponding to the first α-helix of other family members [12]. In our previous study using purified proteins and model membranes, we demonstrated that LC3C induces greater vesicle tethering and membrane fusion than LC3B, and that these differences are largely determined by its N-terminal region [30]. Previous studies identified the N-terminal residues R10 and R11 of LC3B as important for membrane tethering and fusion [25]. However, the corresponding residues in LC3C (R16 and K17) are also positively charged, suggesting that other amino acids contribute to the differences between these proteins. Interestingly, our present experiments with chimeric proteins revealed that the LC3C/B chimera, containing the N-terminal region of LC3C and the ubiquitin-like core of LC3B, formed larger puncta than LC3B, whereas the reciprocal LC3B/C chimera formed smaller puncta, more similar to those of LC3B. These findings suggest a relationship between the membrane-remodeling properties of LC3C characterized *in vitro* and its ability to regulate autophagic vesicle size in a cellular environment. In contrast, the intracellular localization of the chimeric proteins appeared to be primarily determined by their ubiquitin-like cores (Figure 9), suggesting that different structural regions contribute to distinct properties of LC3B and LC3C.

The relationship between the membrane-fusion activity of ATG8 proteins and autophagosome size raises the question of how their N-terminal regions contribute to membrane expansion in cells. ATG2-mediated lipid transfer is considered a major mechanism driving phagophore expansion, with ATG2 transferring lipids from the endoplasmic reticulum to the growing phagophore. This process involves the recruitment and positioning of ATG2 through interactions with WIPI proteins and ATG9A, while ATG9A also facilitates membrane expansion through its lipid scramblase activity [23,50,51]. Our hypothesis is that the membrane-tethering and fusion activities of ATG8 proteins could provide an additional mechanism contributing to the regulation of autophagosome size, acting in a complementary manner to, rather than replacing, ATG2-mediated lipid transfer. Early studies proposed that membrane-anchored ATG8 proteins could promote membrane tethering and fusion through interactions between different membranes *in trans* [24,25]. More recently, Zhang et al [52] identified an additional mechanism in which the N-terminal regions of lipidated LC3 and GABARAP proteins interact with the same membrane to which they are anchored, promoting membrane expansion through interactions *in cis* [53]. Importantly, the *trans* and *cis* mechanisms are not necessarily mutually exclusive, as interactions in trans could facilitate membrane tethering and fusion, whereas interactions in cis could contribute to membrane remodelling and expansion. Interestingly, a recent study using *in situ* cryo-electron tomography identified vesicles associated with the rims of expanding phagophores in yeast cells with impaired Atg2–Atg9 interactions, suggesting that vesicle fusion may contribute to phagophore expansion beyond the initial stages of autophagosome formation [54]. Although the molecular machinery responsible for these fusion events remains to be established, these observations provide a possible cellular context for the membrane tethering and fusion activities of ATG8 proteins. Our present results suggest that the N-terminal region of LC3C could contribute to the regulation of autophagosome size through these membrane-remodelling mechanisms (tethering and fusion), although further experiments are required to understand how this process occurs.

Future studies should investigate the role of LC3C in cellular models expressing sufficient levels of endogenous LC3C to better understand its function under physiological conditions. In addition, a more detailed characterization of the distinct populations of LC3B-and LC3C-positive autophagosomes would help to determine whether they differ in their composition, cargo selectivity, and cellular functions. Finally, identifying the specific sequences within the N-terminal region of LC3C that mediate its membrane tethering and fusion activities in model membranes and in cells could provide further insight into the molecular mechanisms underlying LC3C functional specialization.

## ACKNOWLEDGEMENTS

The authors thank Dr. Sharon Tooze for providing the HexaKO cell line and for sharing protocols for the generation of stable cell lines, and Dr. Harold Jefferies for providing training in this technique. This work was funded in part by the Research Projects Led by Early-Career Researchers of the University of the Basque Country (EHU), Grant EHU-N24/42, and the Basque Government Consolidated Research Groups Program, Grant IT2081-26. U.B. acknowledges the Basque Government for a postdoctoral fellowship (POS_2024_1_0049), and C.A. and D.B. acknowledge the Basque Government for their predoctoral fellowships.

