## Supplementary Figures for "Functional specialization of LC3B and LC3C in autophagosome biogenesis and cargo degradation: relevance of the N-terminal region"

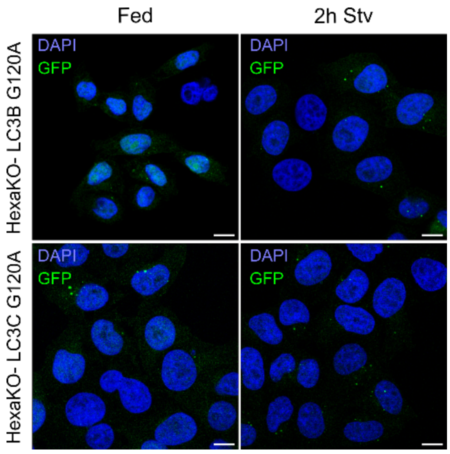


**Supplementary Figure 1. Non-lipidable GFP-LC3B G120A and GFP-LC3C G126A mutants show a predominantly diffuse intracellular distribution without puncta.** Representative fluorescence microscopy images of HexaKO-LC3B G120A and HexaKO-LC3C G126A cells in complete medium (fed) and after 2 h of incubation in EBSS. Scale bar: 10 µm.

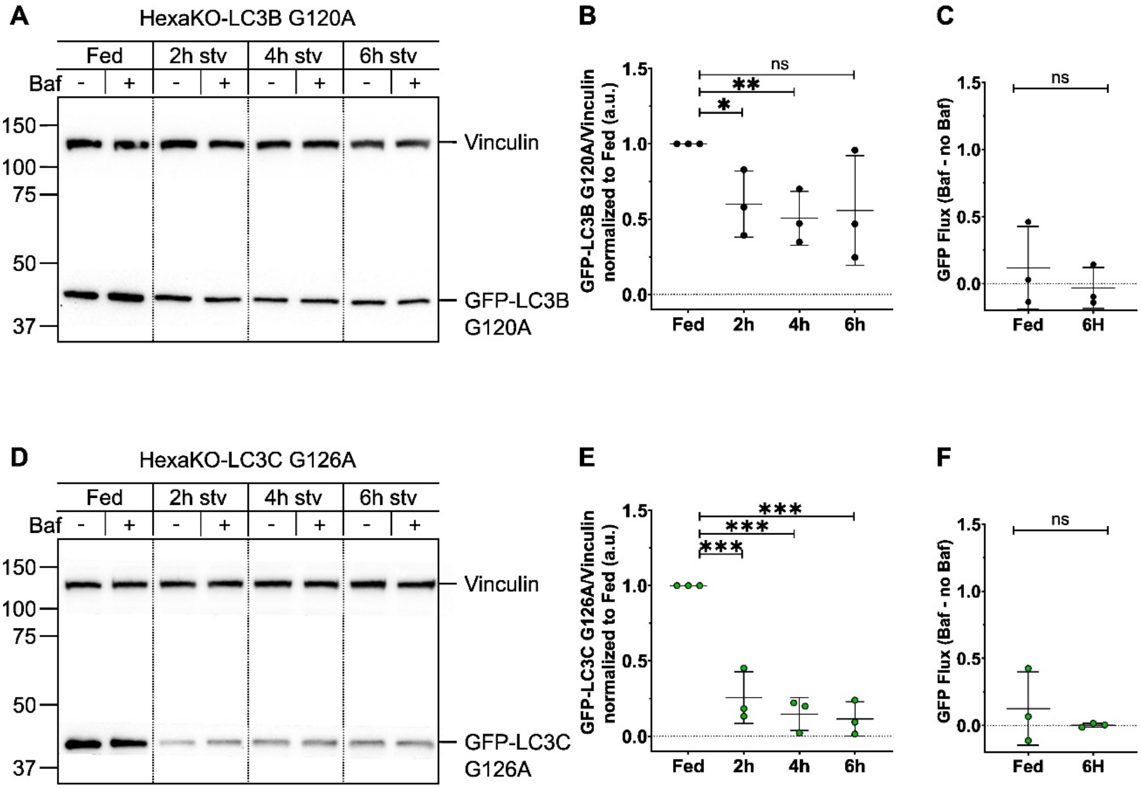


**Supplementary Figure 2. GFP-LC3B G120A and GFP-LC3C G126A degradation under starvation is not prevented by bafilomycin A1.** Representative western blots of HexaKO-LC3B G120A (A) and HexaKO-LC3C G126A (D) cells in complete medium (Fed) and after 2 h, 4 h and 6 h of treatment with EBSS, with or without 100 nM bafilomycin A1. Vinculin was used as a loading control. Quantification of GFP-LC3B G120A (B) and GFP-LC3C G126A (E) levels normalized to Fed. GFP-LC3B G120A (C) and GFP-LC3C G126A (F) flux was calculated by subtracting GFP-LC3 levels in the absence of bafilomycin A1 from those measured in its presence. Values represent the mean ± SD of three independent experiments. Statistical analysis was performed using multiple Student's t-tests. ***p < 0.001; **p < 0.01; *p < 0.05; ns, not significant.

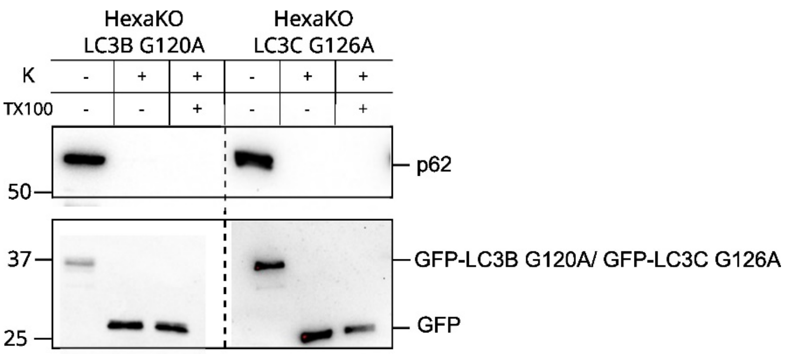


**Supplementary Figure 3. HexaKO-LC3B G120A and HexaKO-LC3C G126A do not form closed autophagosomes.** Representative gel of Proteinase K assay western blots of HexaKO-LC3B G120A and HexaKO-LC3C G126A after 6 h of treatment with EBSS, and 100 nM of bafilomycin A1.

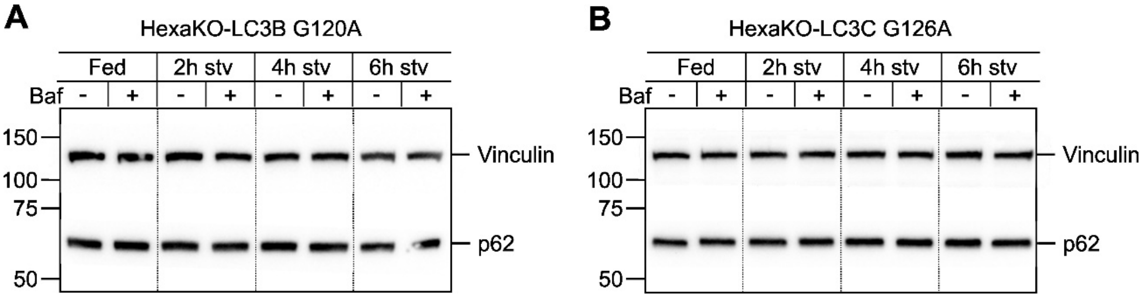


**Supplementary Figure 4. HexaKO-LC3B G120A and HexaKO-LC3C G126A mutants fail to restore p62 degradation.** Representative western blots of HexaKO-LC3B G120A (A) and HexaKO-LC3C G126A (B) in full media (fed) and after 2 h, 4 h and 6 h of treatment with EBSS, with or without 100 nM of bafilomycin A1.

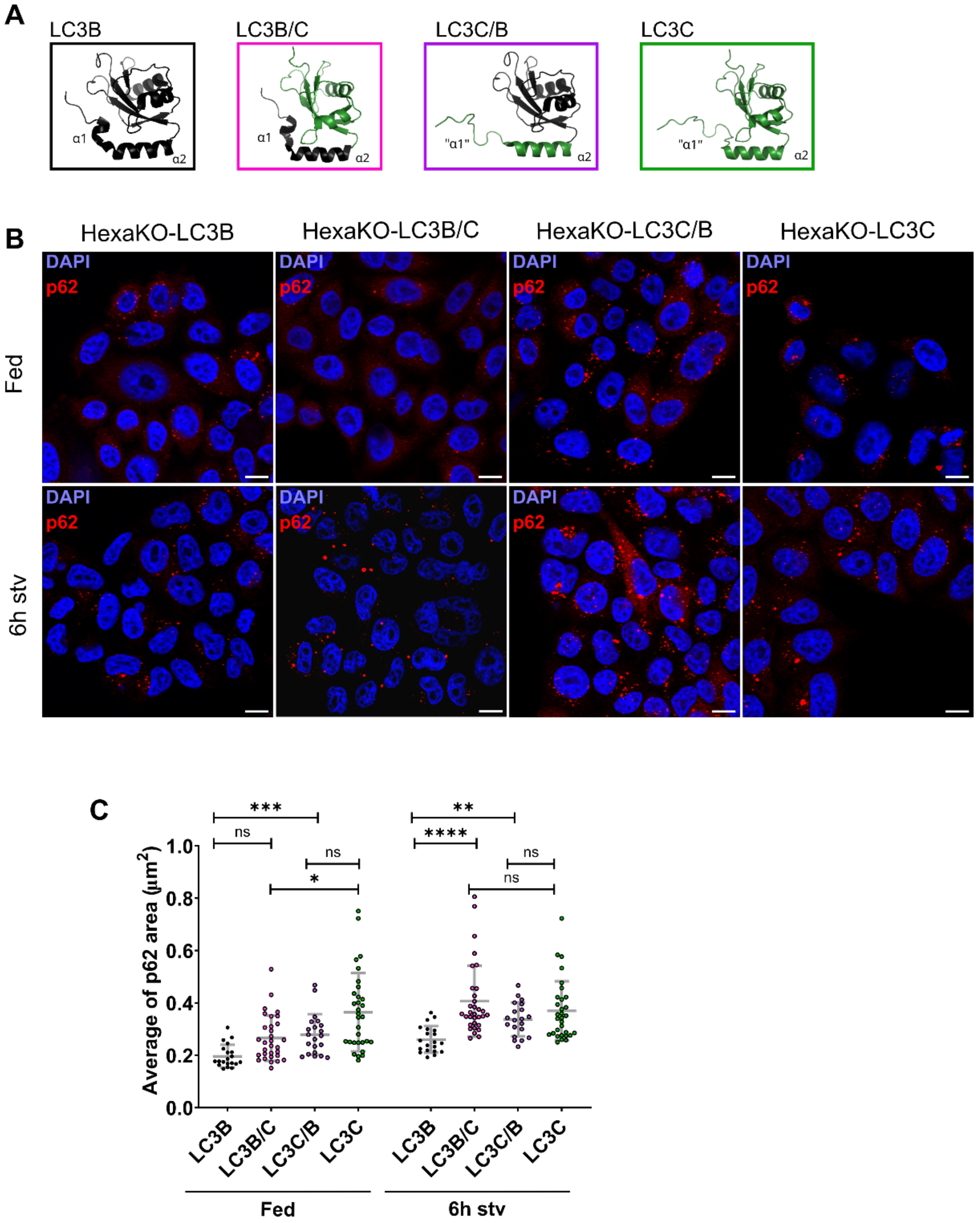


**Supplementary Figure 5. Both the N-terminal region and the ubiquitin-like core of LC3B are required for the efficient reduction of p62 body size.** LC3B (2ZJD) and LC3C (2NCN) structures displayed by PyMOL and the proposed structures of the chimeras LC3B/C (N-terminus of LC3B and ubiquitin-like core of LC3C) and LC3C/B (N-terminus of LC3C and ubiquitin-like core of LC3B) (A). Representative fluorescence microscopy images of HexaKO-LC3B, HexaKO-LC3B/C, HexaKO-LC3C/B and HexaKO-LC3C in full media (fed) and after 6 h of treatment with EBSS (B). Scale bar: 10 µm. Average size (C) of p62-labeled puncta. Average of at least 20 images ± SD. One-way ANOVA significance test: ****p<0.0001; ***p<0.001; ** p<0.01; * p<0.05; ns, non-significant.

**Supplementary Table 1. Vectors and primers used in this study.**

| Vector | Source |
|---|---|
| pHAGE-N-eGFP-LC3B | Dr Christian Behrends (Cluster for Systems Neurology, Munich Germany) |
| pHAGE-N-eGFP-LC3C | Dr Christian Behrends (Cluster for Systems Neurology, Munich, Germany) |
| pHAGE-N-eGFP-LC3B/C | GenScript |
| pHAGE-N-eGFP-LC3C/B | GenScript |
| pHAGE-N-eGFP-LC3B G120A | Dr Christian Behrends (Cluster for Systems Neurology, Munich, Germany) |
| pHAGE-N-eGFP-LC3C G126A | Dr Christian Behrends (Cluster for Systems Neurology, Munich, Germany) |
| pLVX-puro | Dr Sharon Tooze (Francis Crick Institute, London) |
| pLVX-puro-GFP-LC3B | Dr Sharon Tooze ( Francis Crick Institute, London) |
| pLVX-puro-GFP-LC3C | Cloned in this study from pHAGE-N-eGFP-LC3C |
| pLVX-puro-GFP-LC3B/C | Cloned in this study from pHAGE-N-eGFP-LC3B/C |
| pLVX-puro-GFP-LC3C/B | Cloned in this study from pHAGE-N-eGFP-LC3C/B |
| pLVX-puro-GFP-LC3C G126A | Cloned in this study from pLVX-puro-GFP-LC3C |
| pLVX-puro-GFP-LC3B G120A | Cloned in this study from pLVX-puro-GFP-LC3B |
| pLVX-puro-RFP-LC3B | Cloned in this study pHAGE-N-RFP-LC3B |
| **Primers** | **Source** |
| GFP_F: TAAGCAGAATTCGCCACCATGGTGAGCAAGGGC | Merck |
| GFP-LC3C_R: TGCTTAGGATCCCTAGAGAGGATTGCAGGGTCTGTCCTC | Merck |
| GFP-ATG8_R: TGCTTAGGATCCTCGTTATCGAACCACTTTGTACAAGAAAGTTGG | Integrated DNA Technologies (IDT) |
| RFP-LC3B_F: TAAGCAGAATTCATGGCCTCCTCCGAGGACG | Integrated DNA Technologies (IDT) |

| | |
|---|---|
| RFP-LC3B_R: TGCTTAGGATCCTCTGTCACAAGCATGGCTCTCTTCCTG | Integrated DNA Technologies (IDT) |

**Supplementary Table 2. Primary antibodies used for IF**

| Antibody | Dilution | Secondary antibody |
|---|---|---|
| ATG9 (ab108338) | 1:75 | Alexa Fluor 633-conjugated anti-rabbit IgG |
| p62 (sc-28359) | 1:400 | Alexa Fluor 633-conjugated anti-mouse IgG |
| LAMP1 (555798) | 1:50 | Alexa Fluor 633-conjugated anti-mouse IgG |

**Supplementary Video 1 LC3B- and LC3C-positive puncta show limited colocalization.** Stably expressing HexaKO-RFP-LC3B cells were transfected with GFP-LC3C and incubated in EBSS. The video was acquired using a Leica TCS SP5 II confocal microscope, with a frame interval of 30 s, and exported at 10 frames per second. This video is related to Figure 7.
